# Considerations on brain age predictions from repeatedly sampled data across time

**DOI:** 10.1101/2023.03.31.535038

**Authors:** Max Korbmacher, Meng-Yun Wang, Rune Eikeland, Ralph Buchert, Ole A. Andreassen, Thomas Espeseth, Esten Leonardsen, Lars T. Westlye, Ivan I. Maximov, Karsten Specht

## Abstract

**Introduction:** Brain age, the estimation of a person’s age from magnetic resonance imaging (MRI) parameters, has been used as a general indicator of health. The marker requires however further validation for application in clinical contexts. Here, we show how brain age predictions perform for for the same individual at various time points and validate our findings with age-matched healthy controls.

**Methods:** We used densly sampled T1-weighted MRI data from four individuals (from two datasets) to observe how brain age corresponds to age and is influenced by acquision and quality parameters. For validation, we used two cross-sectional datasets. Brain age was predicted by a pre-trained deep learning model.

**Results:** We find small within-subject correlations between age and brain age. We also find evidence for the influence of field strength on brain age which replicated in the cross-sectional validation data, and inconclusive effects of scan quality.

**Conclusion:** The absence of maturation effects for the age range in the presented sample, brain age model-bias (including training age distribution and field strength) and model error are potential reasons for small relationships between age and brain age in longitudinal data. Future brain age models should account for differences in field strength and intra-individual differences.

## Background: What is brain age and what is it good for?

Brain age refers to the estimation of a person’s age from magnetic resonance imaging (MRI) parameters (Franke & Gaser, 2019). This has been done using either neural networks on 3D data (Leonardsen et al., 2022) or tabular data containing region-averaged metrics (Korbmacher et al., 2023; Vidal-Pineiro et al., 2021). Brain age becomes particularly interesting when assuming that lifespan changes in the brain follow normative patterns and that deviations from such patterns might be indicative of disease or disease development (Marquand et al., 2019; Kaufmann et al., 2019). An elevated predicted compared to chronological age in adults may be indicative of psychiatric, neurodegenerative, and neurological disorders (Kaufmann et al., 2019) and poorer health, for example measured by various cardiometabolic risk factors (Beck et al., 2022; Korbmacher et al., 2022). Hence, brain age is a promising developing biomarker of general brain health (Franke & Gaser, 2019).

However, revealing connections between brain age and structural and functional brain architecture is needed to fully understand the biological underpinnings of brain age and its potential clinical implications (Vidal-Pineiro et al., 2021). Furthermore, large cross-sectional samples are often used, which could obscure effects of predictive power of brain age by confounders, in particular, differences in MRI acquisition (Jirsaraie et al., 2022). Hence, contributions of individual differences to brain age estimates require a closer examination. With the aim of assessing the effects of automated MRI scan quality control (QC) metrics on brain age predictions, we used a pre-trained deep neural network model (Leonardsen et al., 2022) to predict brain ages from densly sampled T1-weighted MRI data from three individuals (BBSC1-3) scanned in total N_BBSC_ = 103 times over a one-year interval (Wang et al., 2022), and an independent data set including one individual (FTHP1) scanned N_FTHP_ = 557 times over a three-year interval. We first observed within-subject prediction error and correlations between chronological and predicted age, revealing small, non-significant correlations and larger prediction errors than previously shown in between-subjects analyses. We then tested associations of QC metrics on brain age using linear random intercept models showing potential associations between QC parameters and brain age as well as associations between acquisition parameters and brain age. We validate the findings in cross sectional data and investigate differences in the variability in predictions between longitudinal and cross-sectional datasets.

## Results and Discussion

### Weak correlation between brain age and age

Crude within-subject correlations between age and brain age revealed differing directionalities of slopes across subjects, with only the FTHP1 correlation being statistically significant (*r* = 0.38, 95% CI [0.24; 0.51], *p* < .001; **Figure 1**).

**Figure 1.**
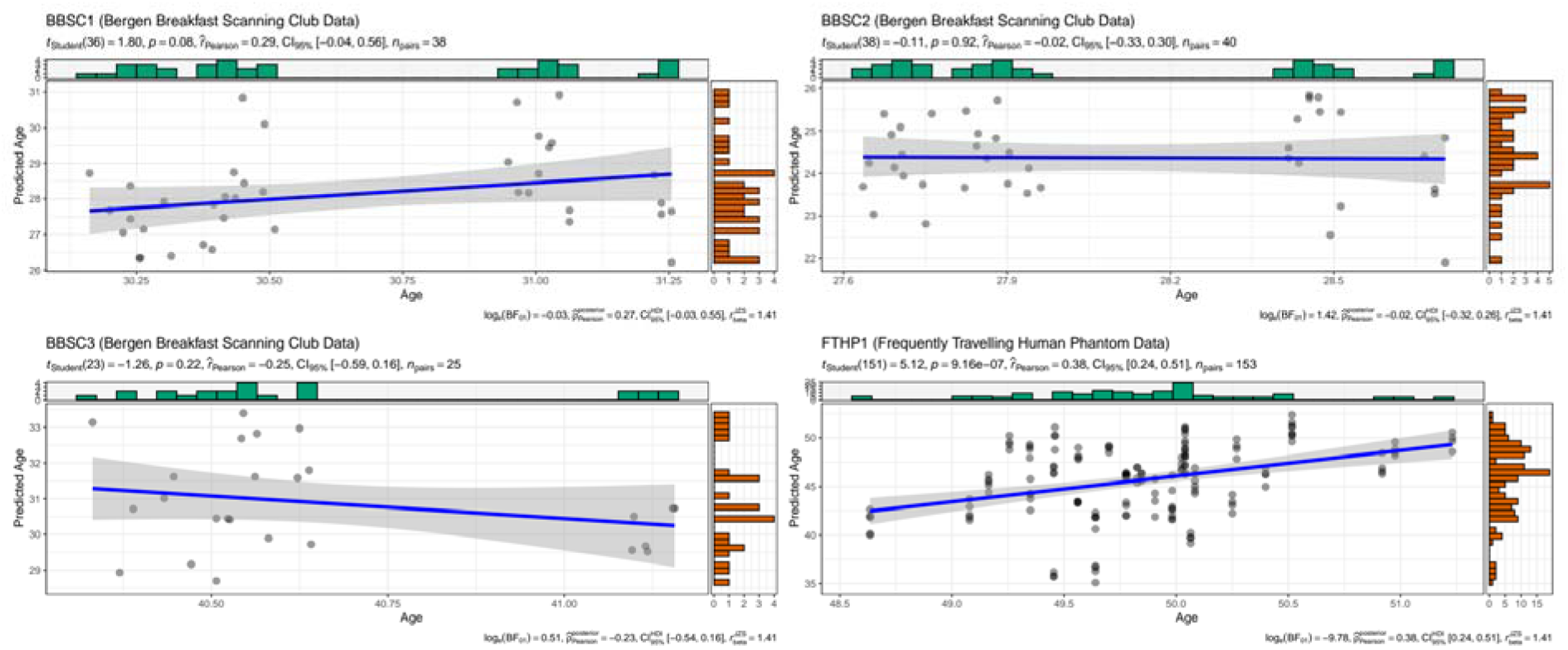
Intra-individual correlations between brain age and chronological age at 3T for BBSC1-3 and FTHP1. Dot colour was grey, with overlapping dots presented as darker.

This is likely due to the small age range and short inter-scan-intervals, as illustrated by differences in model-innate error for the different subjects (**Table 1**) compared to error statistics across age groups (MAE_test_ = 2.47, MAE_external_ = 3.90, as presented in Leonardsen et al., 2022).

**Table 1.** Age, predicted age, brain age gap (BAG), and prediction error by subject and field strength

| Subject | Field Strength | N Observations | Mean Age | SD Age | Mean Prediction | SD Prediction | Mean BAG | SD BAG | MAE | RMSE |
| --- | --- | --- | --- | --- | --- | --- | --- | --- | --- | --- |
| BBSC1 | 3T | 38 | 30.66 | 0.38 | 28.13 | 1.25 | -2.52 | 1.20 | 2.55 | 2.79 |
| BBSC2 | 3T | 40 | 28.09 | 0.38 | 24.37 | 0.95 | -3.72 | 1.03 | 3.72 | 3.85 |
| BBSC3 | 3T | 25 | 40.66 | 0.28 | 30.87 | 1.37 | -9.79 | 1.46 | 9.79 | 9.89 |
| FTHP1 | 3T | 153 | 49.86 | 0.54 | 45.71 | 3.70 | -4.15 | 3.52 | 4.31 | 5.44 |
| FTHP1 | 1.5T | 394 | 49.64 | 0.46 | 48.39 | 2.52 | -1.25 | 2.54 | 2.15 | 2.83 |

The presented data refer to the longitudinal, densely sampled data of few individuals.
BAG = brain age gap, MAE = mean absolute error, RMSE = root mean squared error. BAG is calculated as the difference between predicted age and age.

Additionally, the ages of BBSC1-3 fall into some of the least represented parts of the training data age distribution in the underlying model for the brain age predictions (see Leonardsen et al., 2022) which might contribute to explaining some of the prediction differences beyond model error and intra-individual age range across scanning sessions.

Yet, when using age-matched healthy controls from the cross-sectional TOP and NCNG samples using BBSC and FTHP longitudinal participants’ mean ages ± five years (presented in **Table 1**), correlations between age and brain age estimates were significant for age matches (**Table 2**).

**Table 2.**
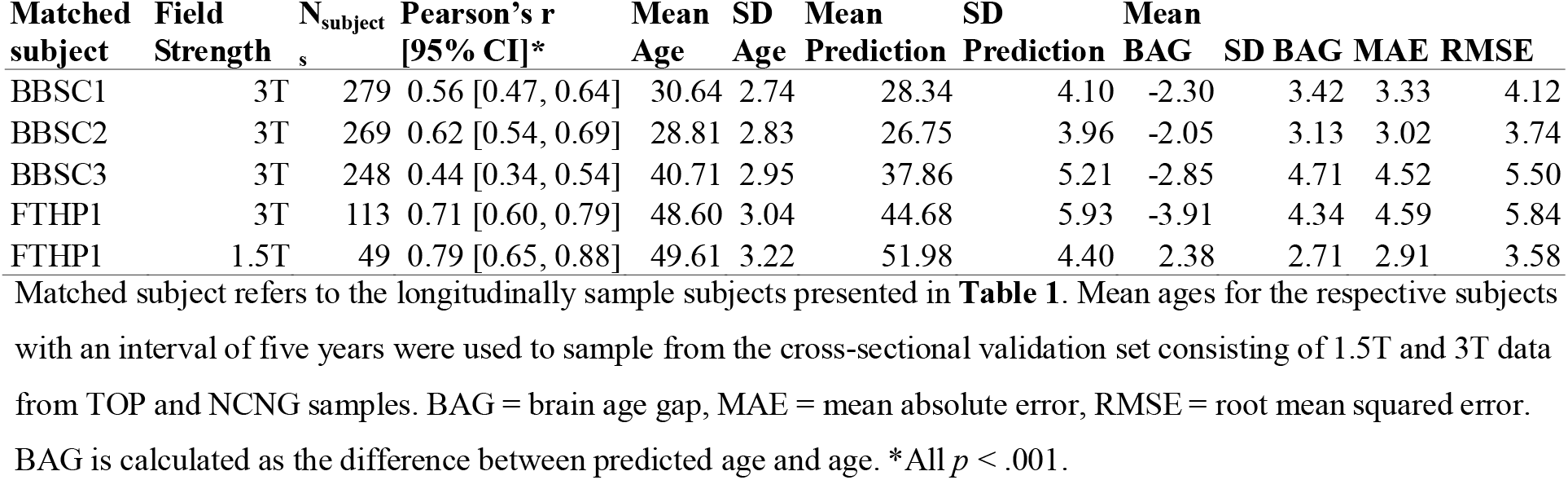
Correlations between age-matching cross-sectional sub-samples’ ages and brain age estimates

| Matched subject | Field Strength | N<sub>subject</sub> | Pearson's r [95% CI]* | Mean Age | SD Age | Mean Prediction | SD Prediction | Mean BAG | SD BAG | MAE | RMSE |
| --- | --- | --- | --- | --- | --- | --- | --- | --- | --- | --- | --- |
| BBSC1 | 3T | 279 | 0.56 [0.47, 0.64] | 30.64 | 2.74 | 28.34 | 4.10 | -2.30 | 3.42 | 3.33 | 4.12 |
| BBSC2 | 3T | 269 | 0.62 [0.54, 0.69] | 28.81 | 2.83 | 26.75 | 3.96 | -2.05 | 3.13 | 3.02 | 3.74 |
| BBSC3 | 3T | 248 | 0.44 [0.34, 0.54] | 40.71 | 2.95 | 37.86 | 5.21 | -2.85 | 4.71 | 4.52 | 5.50 |
| FTHP1 | 3T | 113 | 0.71 [0.60, 0.79] | 48.60 | 3.04 | 44.68 | 5.93 | -3.91 | 4.34 | 4.59 | 5.84 |
| FTHP1 | 1.5T | 49 | 0.79 [0.65, 0.88] | 49.61 | 3.22 | 51.98 | 4.40 | 2.38 | 2.71 | 2.91 | 3.58 |

Interestingly, we also find systematically underestimated brain ages across subjects (**Figure 1)** with the underestimations being stronger for a field strength of 3T than 1.5T for FTHP1 (**Table 1**), and in age-matched cross-sectional data (**Table 2**). While longitudinal brain age predictions were closer related with age at 3T MRI (*r*_*partial*_= 0.38, 95% CI [0.24, 0.51], *p* < .001) than at 1.5T MRI (*r*_*partial*_ = 0.06, 95% CI [-0.04, 0.16], *p* = .239; **Figure 2**), the prediction error was smaller at 1.5T (**Table 1**), with these findings being robust to exclusions of back-to-back repeat scans acquired in the same session without repositioning of the head (**Supplement 1**). When using the out-of-sample test sets TOP and NCNG cross-sectional data, we find highly corresponding relationships between age and brain age at 1.5T (*r* = 0.98, 95% CI [0.97, 0.98], *p* < .001) and 3T (*r* = 0.92, 95% CI [0.91, 0.93], *p* < .001), but higher prediction error at 3T for age matched subjects (**Table 2**) and overall (**Table 3**).

**Table 3.**
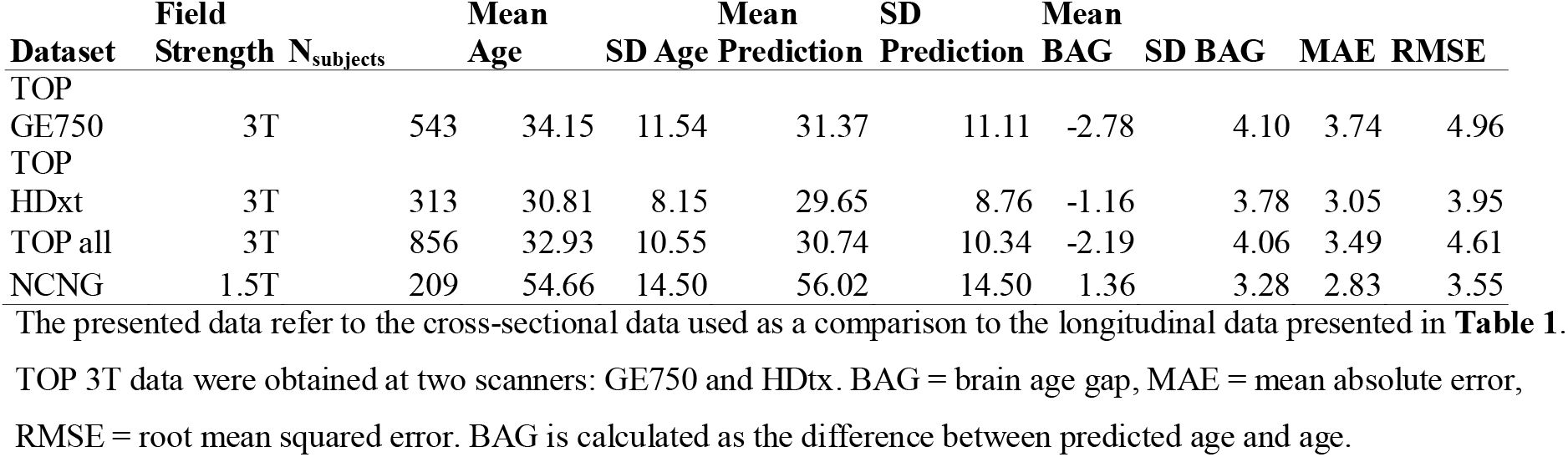
Age, predicted age, brain age gap (BAG), and prediction error by cross-sectional data set and field strength

**Figure 2.**
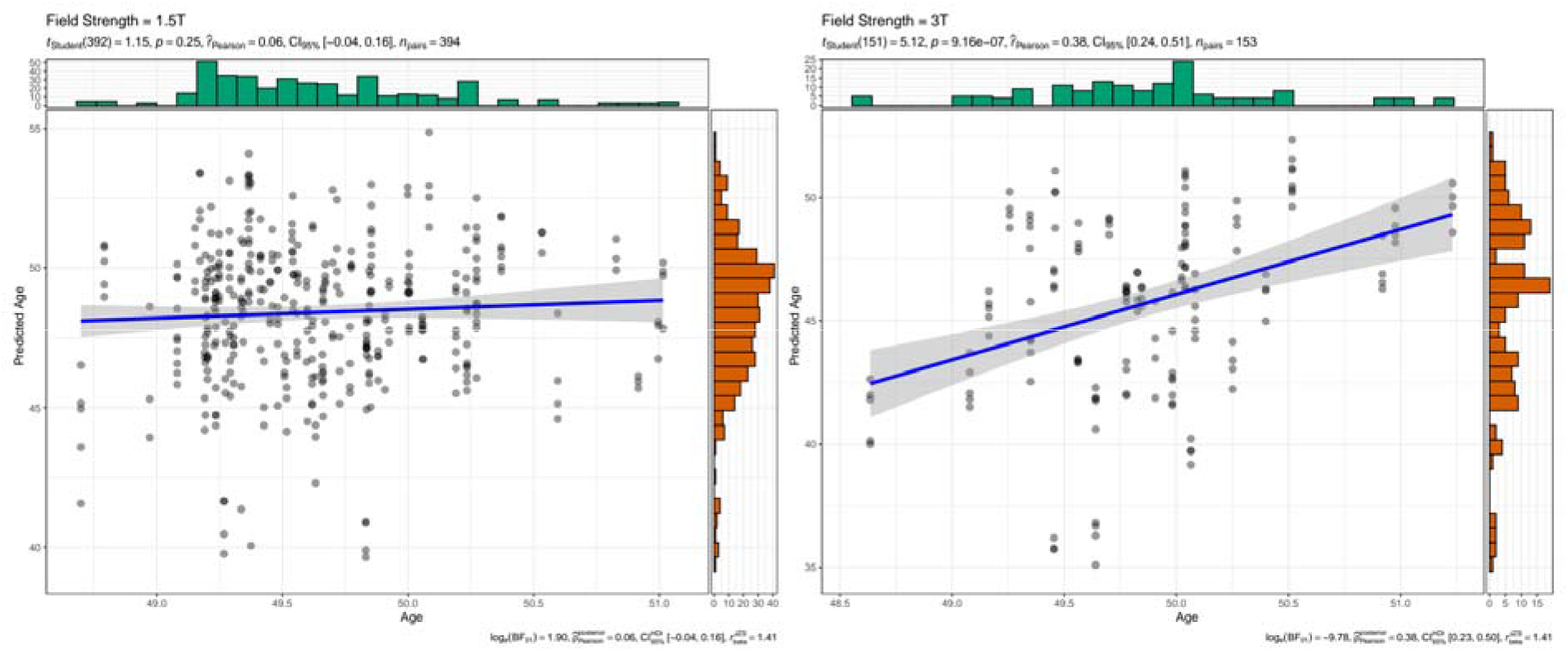
Intra-individual correlations between brain age and chronological age at 1.5T and 3T for FTHP1. Dot colour was grey, with overlapping dots presented darker.

This emphasises the importance of treating predictions for age groups which are underrepresented in the training sample and differences in field strength with care. In that sense, the observed within-subjects variability associated with acquisition- or scanner-specific effects might be used to estimate the minimum size of true within-subject changes (e.g., due to disease) to be detected with a given power. Previous findings outlined the influence of scanner site on brain age predictions and scan quality (Jirsaraie et al., 2022; Leonardsen et al., 2022) indicated by the Euler number (Rosen et al., 2018). Lower quality scans lead to lower prediction errors. We hence hypothesise that there might be additional reasons for inaccuracies in brain age predictions caused by factors beyond the characteristics of the brain age model, in particular scan quality and acquisition parameters.

### Scan quality and acquisition: possible reasons for inaccurate brain age predictions?

We used linear random intercept models at the participant level to examine associations of individual quality control (QC) metrics (see **Figure 3**; **Materials and methods**) and brain age, while controlling for age in BBSC1-3. Entropy-focus criterion (EFC, β_std_ = −0.489, *p*_Holm_ < .001) and the foreground-background energy ratio (FBER, β_std_ = 0.456, *p*_Holm_ < .001) were significant predictors of brain age. In a seperate analysis of FTHP1 (scanned at different sites using different scanning parameters) we included scanner site, field strength, and slice thickness as random factors, rendering none of the QC metrics significant after correcting for multiple testing (*p*_Holm_ = 1).

**Figure 3.**
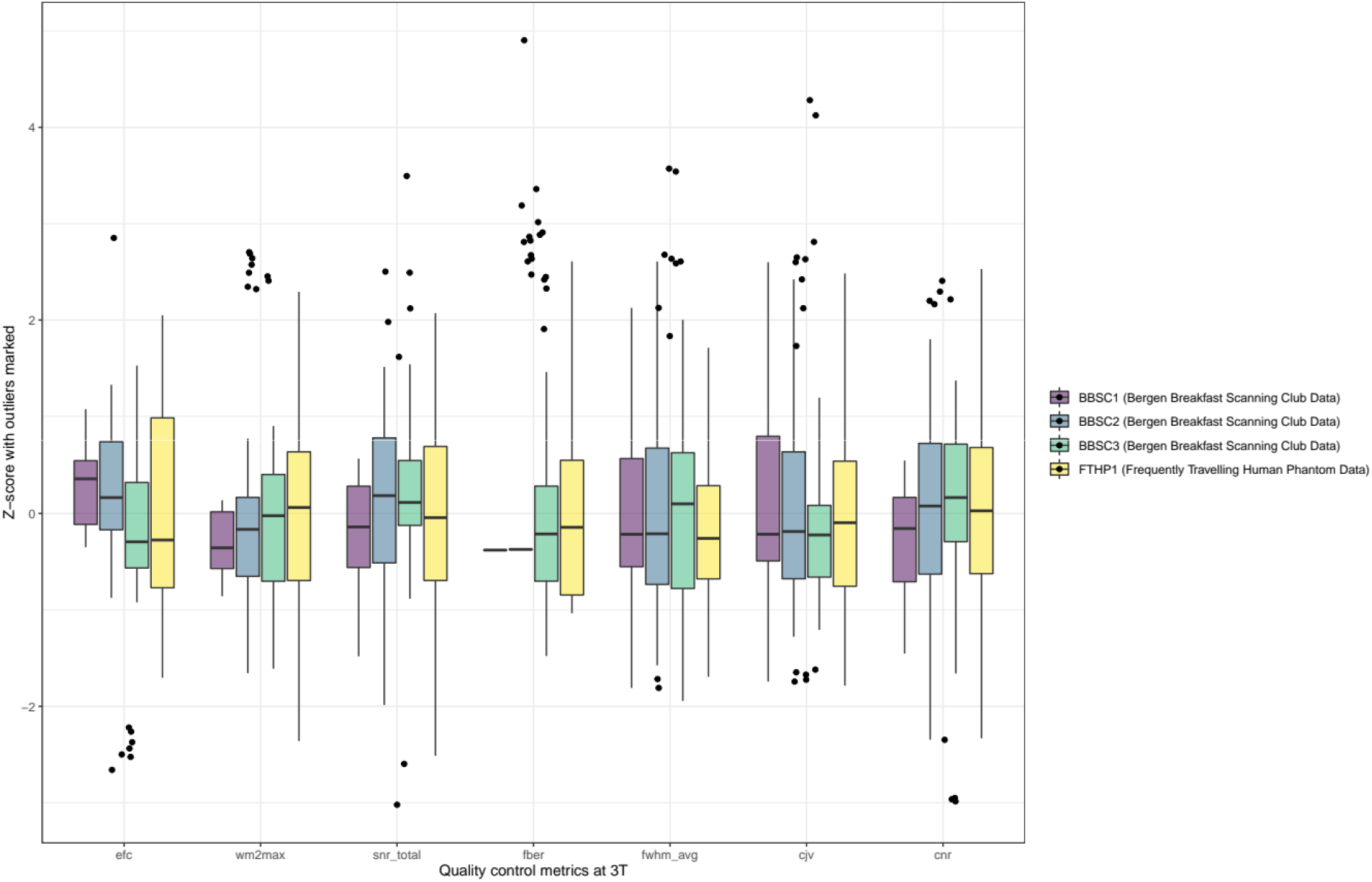
Standardized quality control metrics at 3T per subject. For an overview of scan quality control metrics at 1.5T (only applicable for FTHP1) see **Supplement 2**.

Follow-up analyses in FTHP1 focussed on examining acquisition parameters. We observed individual fixed effects of field strength, manufacturer and slice thickness in one model each, while keeping scanner site and the other acquisition parameters as random effects at the level of the intercept, revealing only significant associations of field strength (β = −1.141, *p*_Holm_ < .001) with brain age.

For validation, we replicate this finding in healthy controls from the TOP and NCNG (see **Materials and Methods** section). BAG was predicted by field strength (β = −2.518, *p* < .001) when controlling for scanner site, with Mean_BAG-1.5T_ = 1.357±3.285 and Mean_BAG-3T_ = −2.19±4.06 using the entire out-of-sample test data. When age-matching FTHP1 and including only the *N* = 162 participants aged 50±5 years (*N* = 49 scanned at 1.5T), the effect of field strength appears stronger (β = −7.40, *p* < .001), with Mean_BAG-1.5T_ = 2.38±2.71 and Mean_BAG-3T_ = −3.92±4.35. In that case, also correlations between age and brain age are stronger at 1.5T compared to 3T (**Table 2**). This was also true when using the entire cross-sectional data (combining TOP and NCNG data), yet with smaller correlational differences when comparing 1.5T (*r* = 0.98, 95% CI [0.97, 0.98], *p* < .001) to 3T (*r* = 0.92, 95% CI [0.91, 0.93], *p* = .004).

While our findings indicate an association between QC parameters EFC and FBER and brain age in all BBSC subjects when controlling for age and constant scanning parameters and scanner site, no QC parameters were significantly associated with brain age after adjustments for multiple comparisons in FTHP1. Based on that, one could speculate that scan quality impacts brain age predictions when particpant ages are sampled from under-represented age groups within the prediction model. For example, Jirsaraie et al. (2022) showed neural networks’ reliability of brain age predictions was lowest add the ends of the age distributions across scanning sites, and predictions were less consistent when image quality was low. Furthermore, QC metrics might be sensitive to individual differences, and vary across scanner sites. FTHP1 results also suggest a strong effect of field strength on brain age. This indicates overall that brain age estimates are potentially dependent on intra-individual variables in addition to acquisition parameters and other scanner site specific covariates. While we cannot generalise from the obtained single-subject results (FTHP1) on field strength, the additional analyses on external datasets support the effect of field strength congruent with Jirsaraie et al.’s (2022) findings of lower prediction errors at 1.5T compared to 3T. This was expressed in our analyses as generally higher brain age estimates at 1.5T compared to 3T, and higher prediction errors at 3T in both cross-sectional and longitudinal data. Finally, we show that prediction error in longitudinal data can be much higher than anticipated from cross sectional estimates, without the presence of mental or physical disorder (see BBSC3 in **Table 1**, compare **Tables 2-3**).

A potential approach for future brain age modelling could be to employ multiple, more specific models which are better tuned to individual differences, developmental trajectories, and scan quality. Such models could for example be trained on data with a smaller age range and a single field strength. Dependent on these parameters, brain age predictions can then be made by a model selected based on the available scan and group the individual belongs to.

## Conclusion

Variability in brain age predictions complicate the metric’s clinical usage, for example, as a (pre-) diagnostic tool. We presented small correlations between age and brain age when repeatedly sampling T1-weighted MRI data from the same individual in a short period of time (1-3 years). Reasons might lay in the absence of maturation effects for the age range in the presented sample, brain age model-bias (including a bimodal or trimodal age training distribution) and model error. While limited, our results suggests an influence of field strength and mixed evidence for scan quality on brain age. Individual differences and the processing of such in the brain age model, might lead to variability in associations between brain age and QC metrics. The presented testing of an established brain age model on multiple single-subject short-timespan retesting data is a stricter test than the usual use-case and does not invalidate MRI group differences. However, intra-individual differences contributing to brain age require further attention in order to advance brain age as a clinical tool.

## Materials and Methods

### Participants

We used two datasets for the analyses which had received ethics approval with all participants consenting formally previously (Opfer et al., 2022; Wang et al., 2022, 2023). The first dataset was the Bergen Breakfast Scanning Club (BBSC) dataset (Wang et al., 2022, 2023), including three male subjects (BBSC2:start-age_BBSC2_ = 27, BBSC1:start-age_BBSC1_ = 30, and BBSC3:start-age_BBSC3_ = 40) who were scanned over the period of circa one year with a summer break in the middle of the scanning period (Wang et al., 2022). This resulted in a total number of N_BBSC_ = 103 scans, relatively equally distributed across subjects (N_BBSC1_ = 38, N_BBSC2_ = 40, N_BBSC3_ = 25). The second dataset was the frequently travelling human phantom (FTHP) MRI dataset (Opfer et al., 2022), including one male subject (FTHP1:start-age_FTHP_ = 48) with 157 imaging sessions at 116 locations, resulting in a total of N_FTHP_ = 557 MRI volumes. Of these, we excluded N = 6 volumes based on errors in the processing pipeline, resulting in a final sample for the main analyses of N_FTHP_ = 551. For quality control (**Supplement 1**), we removed another N_FTHP_ = 25 volumes which were repeat-sequences run at the same scanner and time without changing head position or acquisition parameters, resulting in a final sample for the supplemental analyses of N_FTHP_ = 526.

Finally, as additional validation data, we selected healty controls from two of the cross-sectional out-of-sample test datasets described in Leonardsen et al. (2022): locally collected data (TOP; Tønnesen et al., 2018) and the Norwegian Cognitive NeuroGenetics sample (NCNG; Espeseth et al., 2012), as these provided most MRI scans on healthy controls. Together these datasets include a total of *N* = 209 scans of healthy controls at 1.5T (Mean_age_ = 54.66±15.51), and *N* = 856 scans of healthy controls at 3T (Mean_age_ = 32.93±10.55).

### Image acquisition and preprocessing

T1-weighted volumes of BBSC1-3 were acquired with a spin echo sequence (TE=2.95ms, TR = 6.88ms, FA = 12°, TI = 450, 188 slices, slice thickness = 1mm, in-plane resolution = 1mm×1mm, FOV = 256mm, isotropic voxel size = 1mm^3^) at a 3T GE system with 32-channel head coil (see Wang et al., 2022, 2023). T1-weighted volumes of FTHP1 were acquired at different scanners with various different scanning parameters (see Opfer et al., 2022 or https://www.kaggle.com/datasets/ukeppendorf/frequently-traveling-human-phantom-fthp-dataset). All imaging sites involved in the scanning of FTHP1 were informed that the scan was acquired for the purpose of MRI-based volumetry. Furthermore, all FTHP sites were asked to use acquisition parameters in accordance with the ADNI recommendations for magnetization prepared rapid gradient-echo (MP-RAGE) MRI for volumetric analyses. Thus, the range of FTHP acquisition parameters is representative of MRI-based volumetry in everyday clinical routine at non-academic sites. However, the scan quality might be higher than during average clinical assessments, as only few scans were affected by motion artifacts (relatively young healthy subject). TOP data (Tønnesen et al., 2018) including only healthy controls were acquired at 3T on a GE 3T Signa HDxT (TR = 7.8ms, TE = 2.956ms, FA = 12°; one subset with HNS coil, one subset with 8HRBRAIN coil), and a GE 3T Discovery GE750 (TR = 8.16ms, TE = 3.18ms, FA = 12°). NCNG data (Espeseth et al., 2012) were acquired at a 1.5T Siemens Avanto scanner using two 3D MP-RAGE T1-weighted sequences (TR = 2400 ms, TE = 3.61 ms, TI = 1000 ms, FA = 8°, with 160 sagital slices (1.3 × 1.3 × 1.2 mm)).

Before prediction, the volumes were automatically processed using Freesurfer version 5.3 (Fischl, 2012) and FSL version 6.0 (Jenkinson et al., 2012; Smith et al., 2004), both being widely used open-source software packages (see for overview of advantages and disadvantages compared to other packages: Man et al., 2015) which were validated in clinical and non-clinical samples (Clerx et al., 2015; Fischl, 2012; Jenkinson et al., 2012; Smith et al., 2004). The processing procedure included skull-stripping as part of Freesurfer’s recon-all pipeline, linearly orienting to MNI152 space (6 degrees of freedom) using FSL’s linear registration, and excess border removal. While linear registration in FSL is sensitive to atrophy and high levels of noise (Dadar et al., 2018), this does not apply for the current quality controlled data including only healthy controls. As Freesurfer’s skull stripping algorithm can include errors (Falkovskiy et al., 2016; Waters et al., 2019), the images were manually checked for accuracy. A step-by-step processing tutorial including necessary code can be found at https://github.com/estenhl/pyment-public.

### Brain age estimation

We applied a fully convolutional neural network (Gong et al., 2021; Peng et al., 2021) trained on 53,542 minimally processed magnetic resonance imaging T1-weighted whole-brain images from individuals aged 3-95 collected at a variety of scanning sites (both 1.5 and 3T field strength), (SFCN-reg detailed in Leonardsen et al., 2022) to estimate participants’ ages directly from the MRI using Python v3.9.13. The model was tested in both clinical and non-clinical samples (Leonardsen et al., 2022) and presented high accuracy and test-retest reliability compared to other brain age models (Dörfel et al., 2023).

### Quality control metrics

Quality control (QC) metrics were extracted for each T1-weighted volume by using the automated MRIQC tool version 22.0.6 (Esteban et al., 2017). Of these metrics, we used those which are calculated for the whole brain or volume, being **(1)** noise measures: contrast-to-noise ratio (CNR), signal-to-noise ratio (SNR), coefficient of joint variation of grey and white matter (CJV), **(2)** measures based on information theory entropy-focus criterion (EFC) and foreground-background energy ratio (FBER), **(3)** white-matter to maximum intensity (WM2MAX), and **(4)** other measures: full-width half-maximum (FWHM).

### Statistical analyses

All statistical analyses were conducted using R (v4.1.2). First, correlations of brain age with chronological age and additionally commonly used error metrics for brain age predictions (mean absolute error and root mean squared error) were assessed on a participant level. We further investigated associations between brain age and age in FTHP1 (from the Frequently Travelling Human Phantom dataset) when partialling out scanner site and field strength, as these were expected to influence prediction accuracy.

Further analyses focussed on associations between quality control (QC) metrics and brain age as well as acquisition parameters and brain age. We decided to observe each single independent variable of interest in a dedicated model, as model diagnostics indicated potential multicollinearity in models including multiple QC metrics. Furthermore, random effect models were chosen due to the possibility to account for variances being dependent on different grouping variables, such as ID, scanner site, field strength, and slice thickness.

Hence, linear random intercept models at the particiapant level were used to examine associations of individual QC metrics and brain age, while controlling for age in the BBSC dataset, by running one model for each QC metric. Similarly, for dataset 2, we predicted each QC metric as a fixed effect in addition to the fixed effect of age in a single model. However, we used different random effects, namely, scanner site, field strength, and slice thickness, as dataset 2 contained only FTHP1.

We also examined single individual acquisition parameters in the FTHP dataset (including only one subject FTHP1) as fixed effects in addition to the fixed age effect. Those acquisition parameters of interest were field strength, manufacturer, and slice thickness. Acquisition parameters not used as fixed effects were used as random effect at the level of the intercept in addition to scanner site. All *p*-values were adjusted for multiple testing using Holm correction, marked with *p*_*Holm*_. Standardised β-values (*β*_*std*_) for predictors were used for comparability across β-weights by scaling QC metrics for each subject individually.

Finally, as a validation step, we estimated brain ages for healthy controls in NCNG and TOP datasets and correlated the estimates with age for the entire sample, subjects which were age-matched to the longitudinal, densly sampled individuals mean age ± five years. This provided a baseline understanding for differences in inter and intra subject brain age variability. In a second step, brain age gap (BAG) was examined by field strength and scanner site in the validation sample.

## Acknowledgements

This study was financed by the Research Council of Norway (#276044 and #223273); South-Eastern Norway Regional Health Authority (#2022080); and the European Union’s Horizon2020 Research and Innovation Programme (CoMorMent project; Grant #847776).

