## Supplement for "Considerations on brain age predictions from repeatedly sampled data across time"

### Supplement for "Brain age predictions in longitudinal data reveal the importance of scan quality and field strength"

#### Supplement 1: Analyses on the travelling human phantom data set (data set 2) *excluding* repeat-scans

This section describes the analyses conducted on data set 2, as described in the paper, but without including the N = 30 repeat scans in the analyses.

*Weak relationship between brain age and age (section follow-up)*

Similar to the first data set, in the second dataset, brain age was weakly correlated with chronological age $r_{crude}$ = 0.082, 95% CI [-0.003, 0.166], $p$ = .060 in a subject scanned at multiple sites and varying scanning parameters such as field strength. Brain age predictions were more accurate at 3T ($r_{partial}$ = 0.357, 95% CI [0.208, 0.491], $p$ < .001) than at 1.5T ($r_{partial}$ = 0.066, 95% CI [-0.036, 0.166], $p$ = .204) when holding scanner site constant, as illustrated by a linear model predicting brain age from age ($\beta_{std}$ = 0.166, $p$ <.001), field strength ($\beta_{std}$ = -0.456, $p$ <.001), slice thickness ($\beta_{std}$ = 0.117, $p$ =.002), and scanner ($\beta_{std}$ = -0.125, $p$ =.002). Accordingly, holding field strength constant (ignoring other covariates), strengthened the relationship between age and brain age ($r_{partial}$ = 0.178, 95% CI [0.094, 0.260], $p$ < .001).

*Scan quality: a possible reason for inaccurate brain age predictions?*

Similar to the analysis including repeat scans, none of the QC metrics were significant predictors of brain age ($p_{Holm}$ = 1), and of the acquisition paramters manufacturer, field strength and slice thickness, only field strength was a significant predictors of brain age ($\beta_{\text{std}}$ = -1.156, $p_{\text{Holm}}$ < .001), when using age as fixed effect and site ID,  manufacturer, and slice thickness as random effects.

**Supplement 2: Standardized quality control metrics at 1.5T for FTHP1**

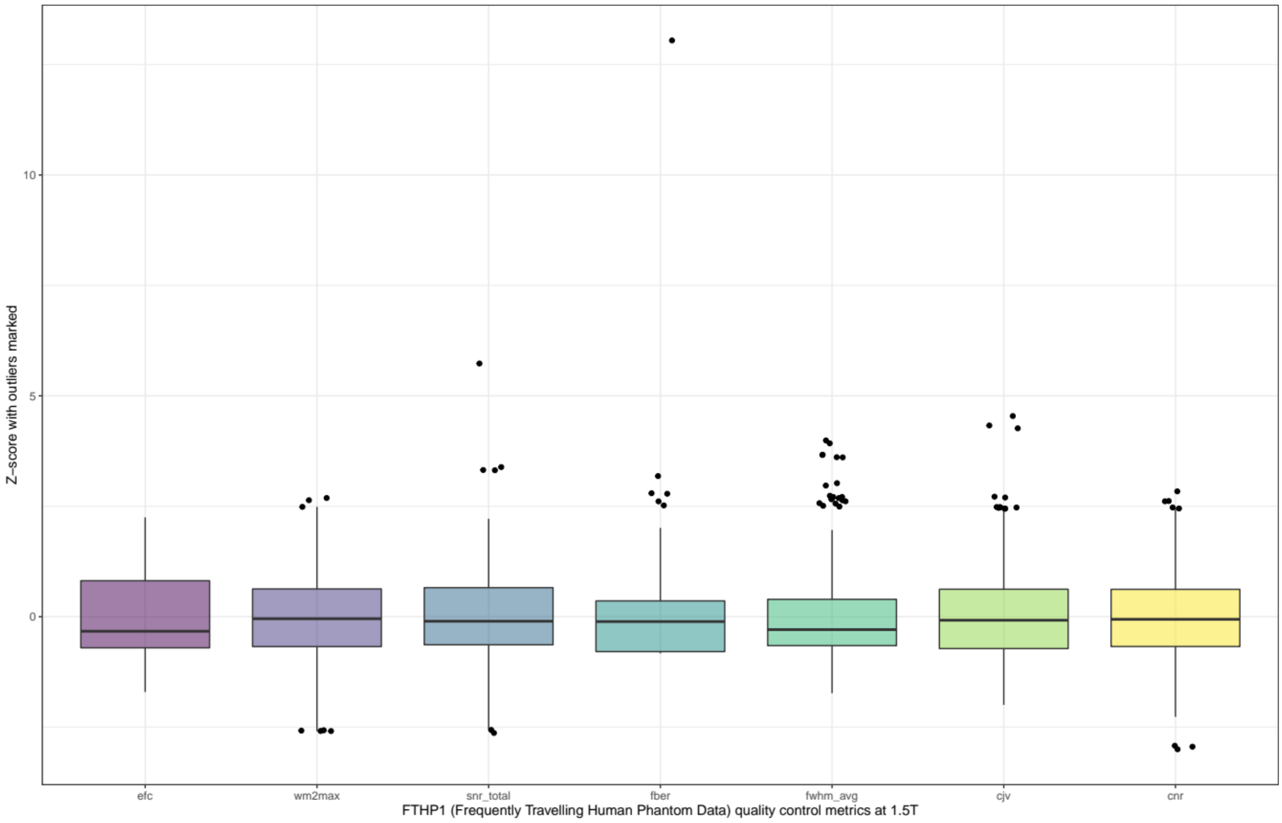


Only for FTHP1 data were collected at 1.5T, hence, qualuty control metrics are only present for this subject.
